# Identification bioactive compounds from marine microorganism and exploration of structure–activity relationships (SARs)

**DOI:** 10.1101/2020.10.23.353169

**Authors:** Mojdeh Dinarvand, Malcolm P. Spain

## Abstract

Marine natural products (MNPs) have become new strong leads for antimicrobial drug discovery and an effective alternative to control drug resistant infections. Herein we report the bioassay guided fractionation of marine extracts from sponges *Lendenfeldia, Ircinia* and *Dysidea* that led us to identify novel compounds with antimicrobial properties. Tertiary amines or quaternary amine salts: anilines **1**, benzylamines **2**, tertiary amines **3** and **4**, and quaternary amine salt **5**, along with three known compounds (**6-8**) were isolated from a crude extract and MeOH eluent marine extracts. The absolute configurations of the new compounds were assigned based on tandem mass spectrometry (MS) analysis. Several of the compounds exhibited potent *in-vitro* antibacterial activity, especially against *Methicillin-resistant Staphylococcus aureus* (MRSA) (MICs from 15.6 to 62.5 micro g/mL). Herein, we also, report structure activity relationships of a diverse range of commercial structurally similar compounds. The structure activity relationships (SARs) results clearly demonstrate that modification of the amines through linear chain length, and inclusion of aromatic rings, modifies the observed antimicrobial activity towards different biological activity. Several commercially available compounds, which are structurally related to the molecules we discovered showed broad spectrum antimicrobial activity against different test pathogens with an MIC50 range of 50 to 0.01 microM. The results of cross-referencing antimicrobial activity and cytotoxicity establish that these compounds are promising potential lead molecules, with a favourable therapeutic index for antimicrobial drug development. Additionally, the SAR studies show that simplified analogues of the isolated compounds with increased bioactivity

## INTRODUCTION

Almost 70 percent of earth’s surface is covered by ocean, representing a huge reserve of natural biological and chemical diversity on our planet.^1^ Marine ecosystems have long been a rich source of bioactive natural products in the search for interesting molecules and novel therapeutic agents.^2–6^ Many interesting and structurally diverse secondary metabolites have been isolated from marine sources over the last 70 years.^7–10^ In addition, the preclinical pharmacology of seventy-five compounds isolated from marine organisms have been reported to have biological activities.^11^ Yet the first ‘drugs from the sea’ were only approved in the early 2000s: the cone snail peptide ziconotide (ω-conotoxin MVIIA) in 2004 to alleviate chronic pain,^12^ and sea squirt metabolite trabectedin in 2007 for treatment of soft-tissue sarcoma.^13^ Marine natural products (MNPs) have displayed exceptional potency and potential as anticancer therapeutics.^14^ Interest in MNPs has continued to grow,^7, 9–10^ spurred in part by the spread of antimicrobial resistant pathogens and the need for new drugs to combat them.^4^

The most prolific marine organisms are sponges,^15^ and the oldest metazoans on earth belong the phylum *Porifera.*^16^ The *Demospongiae* are the most abundant class of *Porifera*, representing 83% of described species,^16–17^ and has the largest number of bioactive compounds.^14^ The genus *Lendenfeldia* are known as a source of sulfated sterols.^18^ The *Lendenfeldia* species metabolites have an anti-HIV, anti-tumor^19^, anti-inflammatory, antifouling^20^ activities but they lack antimicrobial activity.^18^ Secondary metabolites of the genus *Ircinia* and *Dysidea* are prime candidates for further study to unveil their biological metabolites with antibacterial activity.^14, 21–23^

Human pathogens are associated with a variety of moderate to severe infections and the recent rise of multi-drug resistant pathogens makes treatment more difficult. The last two decades have seen the emergence of methicillin-resistant *Staphylococcus aureus* (MRSA) strains resistant even to ‘drugs of last resort’ such as vancomycin,^24^ and *Mycobacterium tuberculosis* resistant to all first-line agents,^25–27^ highlighting the urgent need to find new effective antibiotics with distinct mechanisms of action. Natural products continue to offer a productive source of structural diversity and bioactivity, and are an important source for new drugs.^4–5, 7–8^

In the search for new antimicrobial agents, we screened a set of marine extracts^28^ to determine activity against antibiotic resistant microorganisms using a high-throughput screening (HTS) assay. Fractionation and purification of active components by high-performance liquid chromatography (HPLC), Nuclear magnetic resonance (NMR) and structural elucidation using high resolution and tandem mass spectrometry (MS) led us to a series of potential scaffolds for new, bioactive amine natural products (Figures 1, S20, S21and S24).

**Figures 1.**
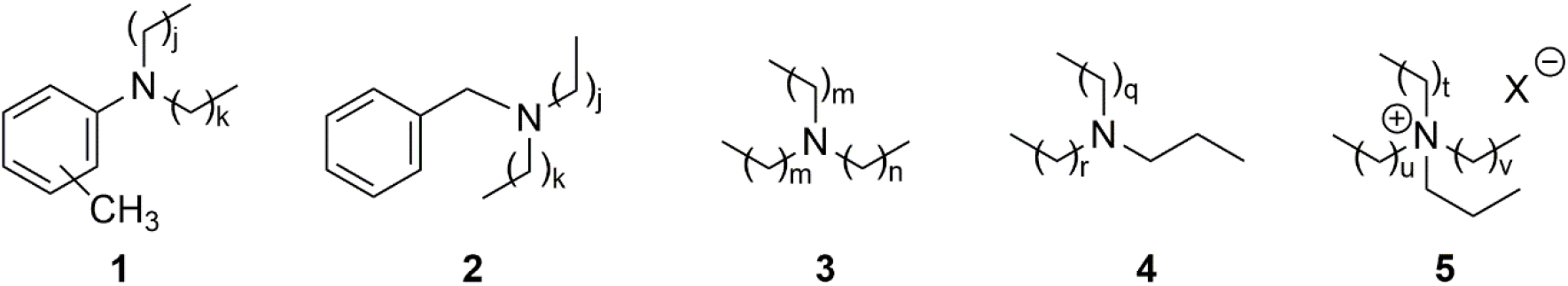
Proposed structures of bioactive amine natural products identified as new compounds in this study; (j + k) = 14; m = 5, n = 9; (q + r) = 20; (t + u + v) = 19; X = unidentified counterion.

## RESULTS AND DISCUSSION

### Identification of active Marine Extracts

To identify marine samples with activity against MRSA, 1434 compounds from the AIMS Bioresources Library^28^ (provided by the Queensland Compound Library,^29^ now called Compounds Australia^30^) were screened in a resazurin cell viability assay. Of the samples tested, 29 inhibited the growth of MRSA by greater than 50% compared to non-treated controls. Minimum inhibitory concentrations (MICs) were determined for the 23 most promising samples, representing extracts and fractions from the phyla Porifera (90%), Echinodermata (5%) and Chordata (5%) (Table 1). The five most active samples showed MICs at 31.3 μg mL^−1^ (all Porifera samples), while another four samples returned MICs of 62.5 μg mL^−1^ (also all Porifera). Cytotoxicity screens against HepG2, HEK 293, A549 and THP-1 cell lines were performed to define the cytotoxicity profile of the most active samples. Pleasingly, all the samples most active against MRSA were also nontoxic to the cell lines tested (Tables 1).

**Table 1.**
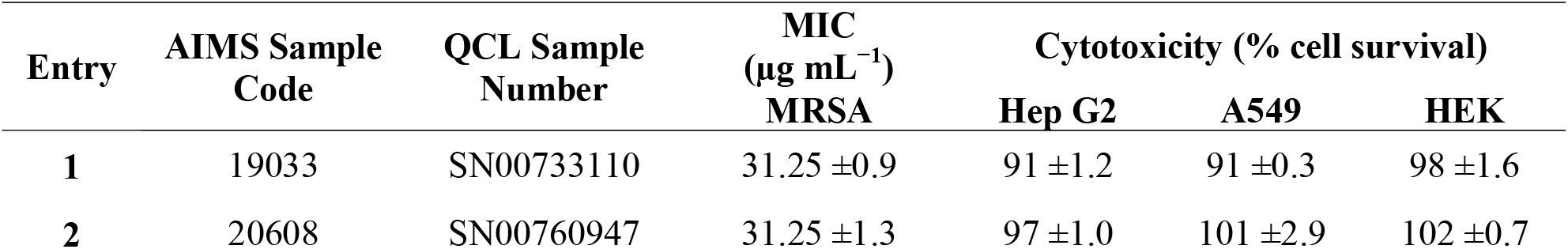
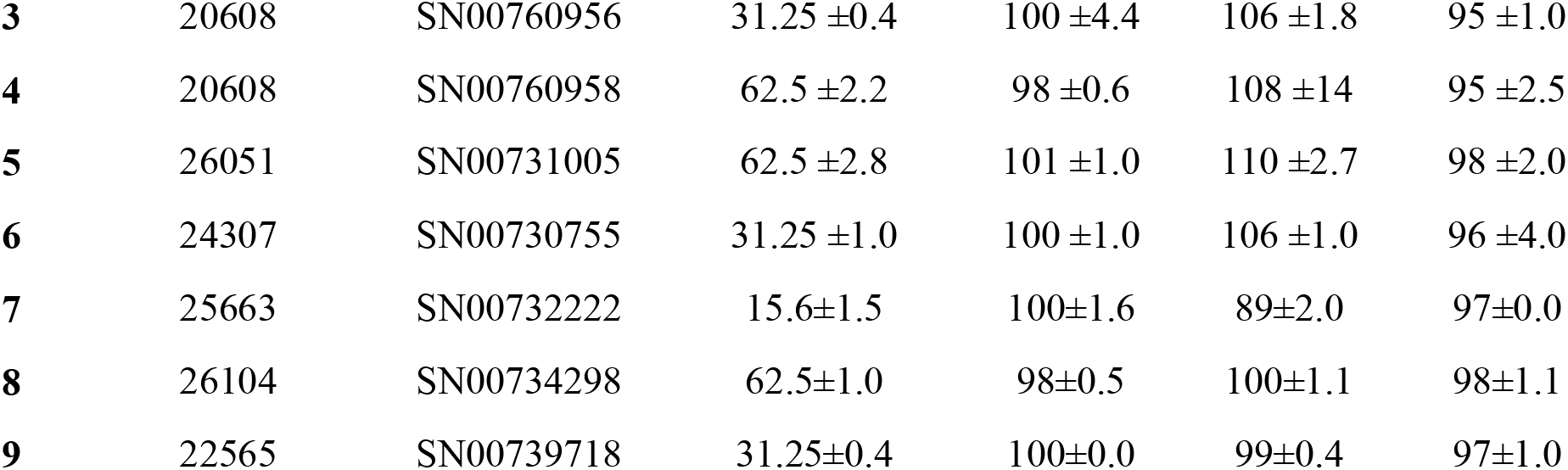
Summary of the Nine Marine Samples Selected for Further Study.

| Entry | AIMS Sample Code | QCL Sample Number | MIC<br>($\mu\text{g mL}^{-1}$)<br>MRSA | Cytotoxicity (% cell survival) | | |
| --- | --- | --- | --- | --- | --- | --- |
|  |  |  |  | Hep G2 | A549 | HEK |
| 1 | 19033 | SN00733110 | $31.25 \pm 0.9$ | $91 \pm 1.2$ | $91 \pm 0.3$ | $98 \pm 1.6$ |
| 2 | 20608 | SN00760947 | $31.25 \pm 1.3$ | $97 \pm 1.0$ | $101 \pm 2.9$ | $102 \pm 0.7$ |
| **3** | 20608 | SN00760956 | 31.25 ±0.4 | 100 ±4.4 | 106 ±1.8 | 95 ±1.0 |
| **4** | 20608 | SN00760958 | 62.5 ±2.2 | 98 ±0.6 | 108 ±1.4 | 95 ±2.5 |
| **5** | 26051 | SN00731005 | 62.5 ±2.8 | 101 ±1.0 | 110 ±2.7 | 98 ±2.0 |
| **6** | 24307 | SN00730755 | 31.25 ±1.0 | 100 ±1.0 | 106 ±1.0 | 96 ±4.0 |
| **7** | 25663 | SN00732222 | 15.6±1.5 | 100±1.6 | 89±2.0 | 97±0.0 |
| **8** | 26104 | SN00734298 | 62.5±1.0 | 98±0.5 | 100±1.1 | 98±1.1 |
| **9** | 22565 | SN00739718 | 31.25±0.4 | 100±0.0 | 99±0.4 | 97±1.0 |

### Isolation and Characterization of Bioactive Compounds

Following the primary screening of the AIMS library and selection of positive hits, HPLC was used to separate and isolate active compounds, guided by bioassays against MRSA. Extracts were fractionated by analytical HPLC (see Experimental section and Supporting Information for further details), and fractions evaluated for bioactivity. Preparative scale HPLC was carried out on each bulk sample to isolate the active component (Table S1, Figures S1–S10), NMR and tandem mass spectrometry (MS/MS) methods used to deduce structures for novel compounds (Table 2 and Supporting Information).^31–32,33^ Insufficient quantities were obtained for positive ion high resolution mass spectrometry (HRMS) analyses.

**Table 2.** Key HRMS Data for Bioactive Samples, and Proposed Structures as Shown in SI.

| Sample | Molecular ion (m/z) | Molecular formula | Proposed structures | Spectra | MS/MS | NMR analyses |
| --- | --- | --- | --- | --- | --- | --- |
| 19033 | 326.37813<br>[M + H]<sup>+</sup> | [C<sub>22</sub>H<sub>48</sub>N]<sup>+</sup><br>Calc. = 326.37802<br>(Δm = 0.11 ppm)<br>RDBE = 0 | **3** | FiguresS11 | Table S2<br>FiguresS12 | - |
| 20608<sup>†</sup> | 332.33115<br>[M + H]<sup>+</sup> | [C<sub>23</sub>H<sub>42</sub>N]<sup>+</sup><br>Calc. = 332.33118<br>(Δm = 0.03 ppm)<br>RDBE = 4<sup>‡</sup> | **1, 2** | FiguresS13 | Table S3<br>FiguresS14 | - |
| 26051 | 368.42508<br>[M + H]<sup>+</sup> | [C<sub>25</sub>H<sub>54</sub>N]<sup>+</sup><br>Calc. = 368.42495<br>(Δm = 0.13 ppm)<br>RDBE = 0 | **4,5** | FiguresS15 | Table S4<br>FiguresS16 | - |
| 25663 | 306.0635<br>[M + Na]<sup>+</sup> | [C<sub>18</sub>H<sub>9</sub>ON<sub>3</sub>]<br>Calc. = 306.0637<br>(Δm = 0.8 ppm)<br>RDBE = 16 | **6** | FiguresS17 | - | Table S5<br>FiguresS20 |
| 25663 | 361.9925<br>[M + H]<sup>+</sup> | [C<sub>18</sub>H<sub>8</sub>BrN<sub>3</sub>O]<br>Calc. = 361.9923<br>(Δm = 0.5 ppm)<br>RDBE = 16 | **7** | Figures18 | - | Table S6<br>FiguresS21 |
| 26104 | 467.117<br>[M + H]<sup>+</sup> | [C<sub>30</sub>H<sub>17</sub>N<sub>2</sub>O<sub>4</sub>]<br>Calc. = 467.118<br>(Δm = 1.9 ppm)<br>RDBE = 0 | **8** | Figures19 | - | Table S7<br>FiguresS22-25 |

<sup>†</sup> The same active species was observed for all five fractions SN00760947, SN00760956, SN00732222, SN00734298 and SN00760958.
<sup>‡</sup> RDBE = ring or double bond equivalents

Active components were isolated and characterised for five of the six extracts shown in Table 1: anilines **1** and benzylamines **2** from the *Lendenfeldia sp.* samples (AIMS Sample Code 20608, Table 1 entries 2-4); tertiary aliphatic amine **3** from the *Dysidea herbacea* extract (AIMS Sample Code 19033, Table 1 entry 1); aliphatic tertiary amine **4** and quaternary amine salt **5** from *Ircinia gigantea* (AIMS Sample Code 26051, Table 1 entry 5); ascididemin **6** and 2-bromoascididemin (2-Bromoleptoclinidinone) **7** from the *Flavobranchia* samples (AIMS Sample Code 25663, Table 1 entries 7); and halisulfate **8** from *Ircinia* (AIMS Sample Code 26104, Table 1 entry 8). Aniline/amines **1**-**5** are all known compounds,^34–36^ however they have not previously been identified as natural products, nor has their antibiotic activity been assessed.

Compounds **6**, **7**^37–38^ and **8**^39–42^ have previously been reported, and in the current study these structures were confirmed by comparison of data with NMR and literature values. In 1994, the first biological activity of compounds **6** was reported as an antitumoral and antineoplastic activity ^43^. Antimycobacterial activity of compounds **6** with wide range of MIC_50_ against different microorganism’s reported ^43, 38,44^. Compounds **7** shows a similar activity profile against these organisms ^38, 45–54^. The inhibitory effects of compounds **6** are not limited to terrestrial microorganisms as it has also shown anti-predatory properties against both puffer and damselfish ^43, 55^.

Compound **8** has previously been identified and isolated from the marine sponges *Theonella swinhoe, Halicondriidea,* and *Coscinoderma mathewsi* and its inhibitory effects on different enzymes reported before ^56–58^. The bioactivity of **8** was first reported in 1998, when it was identified as a thrombin inhibitor ^56^. Antimicrobial activity against *S. aureus*, *C. albicans*, and *B. subtilis* has also been demonstrated ^59^. This compound also inhibited PMA-induced inflammation, phospholipase A2, ATPase activity (with IC_50_ = 8 μM), RNA binding (IC_50_ = 8 μM), serine protease activities (IC_50_ = 14 μM) ^59^ and calcineurin (IC_50_ = 69 μM) ^59–61^.

The helicase-inhibitor (Compound **8**) with biological activity, also detected from other natural sources such as specimens which isolated from marine organisms in Okinawa, Japan, Sorong and Indonesia, has been shown to play an essential role in inhibiting viral replication with IC_50_ values of 3 μM ^59^. Halisulfate analogues with a range of different stereochemistries have been described before reported to have similar activity ^59–61^.

## CONCLUSIONS

Commercial drugs vancomycin and rifampicin remain the main agents for the treatment of invasive MRSA and *M. tuberculosis* diseases, respectively. Yet the number of vancomycin-resistant *S. aureus* (VRSA) and rifampicin-resistant *M. tuberculosis* strains is on the rise. The emergence of antibiotic resistance highlights the need for novel, effective antibacterial agents which circumvent traditional resistance mechanisms.

Thus we assayed 1434 extracts from the AIMS Bioresources Library^28^ against MRSA, finding five samples that have a promising combination of high antibacterial activity and low toxicity to mammalian cells: AIMS Sample Codes 20608, 26051, 25663, 26104 and 19033 (Table 1). Samples of three extracts (20608, 26051 and 19033) were subjected to HPLC purification and bioassay guided fractionation, enabling bioactive components to be isolated in low yield («1 mg). Then high-resolution MS and tandem MS analysis was used to decipher structures (Table 2, Figures1). The proposed structures are all tertiary amines or quaternary amine salts: anilines **1** and benzylamines **2** (from *Lendenfeldia* sample number 20608), aliphatic amine **3** (from *Dysidea herbacea* sample number 19033), aliphatic tertiary amines **4** and quaternary amine salt **5** (from *Ircinia sp.* sample number 26051). The proposed structures from samples 25663 and 26104 are ascididemin **6** and 2-bromoascididemin **7** (from *Flavobranchia* sample number 25663), and halisulfate **8** (from *Ircinia* sample number 26104),

The compounds uncovered in this study add to the growing arsenal of antimicrobial agents from the sea,^3–4^ and offer interesting new avenues in the quest for new, effective agents to combat the growing scourge of multidrug resistant bacteria.

## EXPERIMENTAL SECTION

### General

Chemical reagents were purchased from BDH Chemicals and Sigma Aldrich (Castle Hill, Sydney, Australia) and used as supplied unless otherwise indicated.

### Natural Product Library

Natural product extracts were provided by the Australian Institute of Marine Science (AIMS), Townsville, Queensland as part of the AIMS Bioresources Library,28 via the Queensland Compound Library,29 (now called Compounds Australia30). Crude extracts had been partially fractionated by AIMS/ QCL to generate a library of 1434 samples, supplied in DMSO (100%) solution and stored at −80 °C. Original concentrations as provided were 5 mg mL^−1^. Stock solutions were made by diluting these samples by a factor of 1:10 in dH_2_O and stored at −80 °C.

### Bacterial Inhibition Assays

For screening of the AIMS library, each test sample (10 μL) was dispensed into a separate well of a 96 well microtiter plates (final sample concentration 0.5 mg mL^−1^) using sterile dH_2_O. For determination of MIC, extracts (250 to 0.5 μg mL^−1^) or synthesized compounds (100 to 0.0002 μM) were serially diluted in microtiter plates. Bacterial suspension (90 μL, OD600nm 0.001) was added to each well and plates were incubated at 37°C for either 18 hours (MRSA, P. aeruginosa PAO1, E. coli EC958) or 7 days for M. tuberculosis H37Rv as described previously.49-50 Resazurin (10 μL; 0.05% w/v) was added and plates were incubated for 3 h or 24 h (*M. tuberculosis*) at 37 °C. The inhibitory activity was calculated by visual determination of colour change within wells or detection of fluorescence at 590 nm using a FLUOstar Omega microplate reader (BMG Labtech, Germany). Percentage survival was calculated in comparison to the average of untreated control wells after normalising for background readings.

### Evaluating Toxicity of AIMS Extract Library

Human alveolar epithelial cells (A549), Madin-Darby canine kidney epithelial cells (MDCK),^63^ human leukaemia cells (THP-1),^64^ human hepatocellular carcinoma cells (Hep-G2),^65^ and human embryonic kidney cells 293 (HEK293)^66^ were grown and differentiated in complete RPMI (Roswell Park Memorial Institute Medium) and DMEM (Dulbecco’s Modified Eagle’s medium) tissue culture media (RPMIc and DMEMc). To determine toxicity of the AIMS extract library, 2 × 10^5^ of each cell type were added to a 96-well plate and left for 48 h at 37 °C to adhere. Extract samples at a final concentration of 0.5 mg mL^−1^ were added to the wells, then incubated for 7 days in a humidified 5% CO_2_ incubator at 37 °C. Then resazurin (10 μL of 0.05% w/v) was added and after 4 h, fluorescence measured as described previously. Cell viability was calculated as percentage fluorescence relative to untreated cells.

#### Purification of Natural Products from Extracts and Structure Elucidation

##### High Performance Liquid Chromatography (HPLC) Purification

Samples were separated using analytical (Waters 2695 Alliance pump with Waters 2996 PDA, Sunfire reversed-phase column, and WFIII fraction collector) and preparative (Waters 600 HPLC pump, Phenomenex reversed-phase column, Waters 2487 UV detector and WFIII fraction collector) HPLC systems with UV detectors at 254 and 280 nm, employing a gradient of solvents A (dH_2_O) and B (acetonitrile) with trifluoroacetic acid (0.1%). Extract mixtures were kept at 4 °C until injection, then extract sample (100 μL) was injected onto an analytical Waters X-bridge C18 100 Å (4.6 × 250 mm, 5 μm) reversed-phase column on the same analytical HPLC system described above. The mobile phase was obtained using binary gradients of solvents A and B at a flow rate of 1 ml min^−1^ at 30 °C over 80 min. Fractions, separated every 60 s, were collected. Purified fractions were flash-frozen in liquid nitrogen then freeze-dried overnight. The resulting fractionated extracts were re-suspended in DMSO and antibacterial activity versus MRSA was determined as described above.

Fractions identified as active against MRSA were further purified on the preparative HPLC unit described above, using a Phenomenex C18 100 Å (250 × 21.2 mm, 10 μm) reversed-phase column with UV detection at 254 and 280 nm, 7 mL min^−1^ flow rate with water/acetonitrile gradient containing 0.1% trifluoroacetic acid.

The gradient for AIMS extracts 19033, 20608, 26104, 25641and 26051 was 0% B initially, increased to 40% B over 20 min, then to 100% over 40 min, held at 100% for 10 min. For AIMS extracts 25663 gradient was 0% B initially, increased to 40% B over 60 min, then to 100% over 30 min, held at 100% for 10 min. Compounds thus purified were evaluated for biological activity and analysed by MS to determine potential structures for the bioactive components.

#### Identification and Structure Elucidation

##### Identification and elucidation of purified compounds from sample numbers 19033, 20608 and 26051

Purified compounds were identified and characterised using MS and NMR. High resolution ESI mass spectra (HRMS) were recorded on a Bruker Apex Qe 7T Fourier Transform ion cyclotron resonance mass spectrometer with an Apollo II ESI MTP ion source with samples (in CH_3_CN:H_2_O 1:1) infused using a Cole Palmer syringe pump at 180 μL h^−1^. Where required, low resolution ESI tandem MS was performed on a Bruker amaZon SL ion trap via syringe infusion or by injection into a constant flow stream with a rheodyne valve and an Alltech HPLC pump (mobile phase methanol, flow rate 0.3 mL min^−1^) connected to an Apollo II ESI MTP ion source in positive ion mode. Tandem mass spectra of the [M+H]^+^ parent ion were obtained manually up to MS^5^ (depending on sensitivity). Spectra were acquired in positive ion mode using a 1–4 Da isolation window, with the excitation amplitude manually optimized for each spectrum to have the selected mass at ~10% of the height of the largest fragment. Data analysis was performed for both high resolution MS and low resolution tandem MS data using Bruker Data Analysis 4.0 with smart formula assuming C, H, N, O, Na (0-1), mass error <2 ppm, C:H ratio 3 maximum, even electron (or both for tandem MS data). The results of high resolution MS data analysis were further refined manually by comparing isotopic fine structures of simulations where possible (resolving power > 200,000) to further eliminate potential formulae within the 2ppm mass error window (particularly ^15^N, ^18^O, ^2^H, ^13^C and ^13^C_2_ isotopes and confirm no ^34^S presence).

### Identification and elucidation of fractions 44 from sample number 25663 (Compound 6)

^1^H NMR (500 MHz, DMSO-*d*_6_): δ 9.20 (d, *J*=5.6, 1H, H-21), 9.10 (dd, *J*=4.5, 1.7, 1H, H-17), 9.00 (dd, *J*= 8.0, 1.1, 1H, H-6), 8.92 (d, *J*= 5.6, 1H, H-19), 8.63 (dd, *J*=7.9, 1.7, 1H, H-15), 8.44 (d, *J*= 8.0, 1H, H-3), 8.09 (ddd, *J*= 8.0, 7.2, 1.1, 1H, H-2), 8.02 (ddd, *J*= 8.0, 7.2, 1.1, 1H, H-1), 7.80 (dd, *J*=7.9, 4.5, 1H, H-16). **HRMS** (ESI): *m/z* 306.0635; the molecular formula C_18_H_9_ON_3_ gives an expected molecular [M+Na]^+^ ion at 306.0637 (err 0.8 ppm). This molecular formula gives a rdbe of 16, requiring many rings or double bonds. The NMR and mass data are in agreement with those in the literature for the known compound ascididemin ^38, 67^.

### Identification and elucidation of fractions 58 from sample number 25663 (Compound 7)

^1^H NMR (500 MHz, DMSO-d_6_): δ 9.25 (d, J=5.3, 1H, H-21), 9.13 (d, J=4.3, 1H, H-17), 8.98 (d, J= 8.8, 1H, H-6), 8.93 (m, 1H, H-19), 8.68 (s, 1H, H-15), 8.64 (d, J= 7.6, 1H, H-3), 8.19 (d, J= 8.8 1H, H-1), 7.81 (dd, J=4.3, 7.6, 1H, H-16). HRMS (ESI): m/z 361.9925; the molecular formula C18H8BrN3O would give an expected [M+H]^+^ ion at 361.9923 (err 0.5 ppm). This molecular formula gives a rdbe of 16, requiring many rings or double bonds. The NMR and mass data are in agreement with those in the literature for the known compound 2-bromoascididemin ^68^.

### Identification and elucidation of purified compounds from sample number 26104 (Compound 8)

^1^H NMR (500 MHz, DMSO-d_6_): δ 8.47 (br s, 1H, H-8), 8.46 (br s, 1H, H-7), 6.54 (d, J= 8.4, 1H, H-6), 6.44 (d, J= 3.2, 1H, H-3), 6.35 (dd, J=3.2, 8.4, 1H, H-5), 5.29 (m, 1H, H-25), 5.22 (t, J= 7.3, 1H, H-10), 4.19 (d, J= 11.4, 1H, H-18), 3.12 (d, J=7.3, 1H, H-9), 2.19 (m, 1H, H-21), 2.13 (m, 1H, H-16), 1.95 (t, J= 7.4, 1H, H-13), 1.91 (m, 1H, H-24), 1.8 (m, 1H, H-24′), 1.66 (d, J= 13.1, 1H, H-27), 1.62 (s, 3H, H-12), 1.6 (s, 3H, H-31), 1.42 (m, 1H, H-14), 1.41 (m, 1H, H-28), 1.37 (m, 1H, H-29), 1.34 (m, 1H, H-20), 1.21 (m, 1H, H-15), 1.13 (m, 1H, H-23), 1.12 (m, 1H, H-29′), 1.06 (m, 1H, H-20′), 1.03 (m, 1H, H-27′), 1.03 (m, 1H, H-15′), 0.84 (s, 3H, H-33), 0.83 (s, 3H, H-34), 0.78 (d, J= 6.9, 1H, H-26), 0.66 (s, 3H, H-32). LRMS (APCI): m/z 467.3, [M+H]^+^; HRMS (APCI): m/z 467.1, [M+H]^+^; the molecular of formula C_30_H_17_N_2_O_4_ calculated at 467.118, err 1.9 ppm. This results in rdbe calculation of 24 indicating a high level of unsaturation and rings. The NMR and mass data are in agreement with those in the literature for the known compound halisulfate ^69–72^.

## ASSOCIATED CONTENT

### Supporting Information

Bioactivity and toxicity screening data, HPLC fractionation and purification protocols, plus mass spectrometry data (HRMS spectra, tables of daughter ions, and proposed fragmentation pathways) for natural products and synthetic compounds.

## AUTHOR INFORMATION

### Author Contribution Statement

MD conceived and designed the experiments; performed the experiments; analyzed the data; MD wrote the paper.

### Notes

The authors declare no competing financial interest

## Acknowledgement

We thank Dr John Merlino (Concord Hospital, Sydney) for provision of the MRSA. This work was supported by the National Health and Medical Research Council (NHMRC) Project APP1084266, the NHMRC Centre of Research Excellence in Tuberculosis Control (APP1043225), and the University of Sydney. MD was supported by an International Postgraduate Research Scholarship (IPRS) and Australian Postgraduate Award (APA) from the Australian Government.

## ABBREVIATIONS

A549: human alveolar epithelial cells
AIMS: Australian Institute for Marine Science
DMEM: Dulbecco’s Modified Eagle’s medium
HEK293: human embryonic kidney cells 293
HPLC: high-performance liquid chromatography
HTS: high-throughput screening
MDCK: Madin-Darby canine kidney epithelial cells
MIC: minimum inhibitory concentration
MRSA: methicillin resistant *Staphylococcus aureus*
MS: mass spectrometry
MS/MS: tandem mass spectrometry
MTC: minimum toxic concentration
NMR: nuclear magnetic resonance
RPMI: Roswell Park Memorial Institute Medium
THP-1: human leukaemia cells
Hep-G2: human hepatocellular carcinoma cells
VRSA: vancomycin-resistant *Staphylococcus aureus*.

## REFERENCES

1. El-Demerdash A, Atanasov AG, Horbanczuk OK, Tammam MA, Abdel-Mogib M, Hooper JNA, Sekeroglu N, Al-Mourabit A, Kijjoa A. (2019). Chemical Diversity and Biological Activities of Marine Sponges of the Genus Suberea: A Systematic Review. Marine drugs 17:115.

2. Molinski TF, Dalisay DS, Lievens SL, Saludes JP. (2008). Drug development from marine natural products. Nature Reviews Drug Discovery 8:69–85.

3. Hughes CC, Fenical W. (2010). Antibacterials from the Sea. Chemistry – A European Journal 16:12512–12525.

4. Indraningrat A, Smidt H, Sipkema D. (2016). Bioprospecting Sponge-Associated Microbes for Antimicrobial Compounds. Marine Drugs 14:87.

5. Mehbub MF, Perkins MV, Zhang W, Franco CMM. (2016). New marine natural products from sponges (Porifera) of the order Dictyoceratida (2001 to 2012); a promising source for drug discovery, exploration and future prospects. Biotechnol Adv 34:473–491.

6. Schroeder G, Bates SS, La Barre S. (2018). Bioactive Marine Molecules and Derivatives with Biopharmaceutical Potential. In Barre SL, Bates SS (ed), Blue Biotechnology doi:doi:10.1002/9783527801718.ch19.

7. Newman DJ, Cragg GM. (2016). Natural Products as Sources of New Drugs from 1981 to 2014. Journal of Natural Products 79:629–661.

8. Pye CR, Bertin MJ, Lokey RS, Gerwick WH, Linington RG. (2017). Retrospective analysis of natural products provides insights for future discovery trends. Proceedings of the National Academy of Sciences of the United States of America 114:5601–5606.

9. Blunt JW, Copp BR, Keyzers RA, Munro MHG, Prinsep MR. (2017). Marine natural products. Natural Product Reports 34:235–294.

10. Blunt JW, Carroll AR, Copp BR, Davis RA, Keyzers RA, Prinsep MR. (2018). Marine natural products. Natural Product Reports 35:8–53.

11. Mayer AMS, Rodríguez AD, Taglialatela-Scafati O, Fusetani N. (2017). Marine Pharmacology in 2012-2013: Marine Compounds with Antibacterial, Antidiabetic, Antifungal, Anti-Inflammatory, Antiprotozoal, Antituberculosis, and Antiviral Activities; Affecting the Immune and Nervous Systems, and Other Miscellaneous Mechanisms of Action. Marine drugs 15:273.

12. McGivern JG. (2007). Ziconotide: a review of its pharmacology and use in the treatment of pain. Neuropsychiatric disease and treatment 3:69–85.

13. Gordon EM, Sankhala KK, Chawla N, Chawla SP. (2016). Trabectedin for Soft Tissue Sarcoma: Current Status and Future Perspectives. Advances in Therapy 33:1055–1071.

14. El-Damhougy KA HAE-N, Hassan AH Ibrahim, Mansour AE Bashar and Fekry M Abou Senna. (2017). Biological activities of some marine sponge extracts from Aqaba Gulf, Red Sea, Egypt. International Journal of Fisheries and Aquatic Studies 5.

15. Indraningrat AA, Smidt H, Sipkema D. (2016). Bioprospecting Sponge-Associated Microbes for Antimicrobial Compounds. Mar Drugs 14.

16. Matobole RM, van Zyl LJ, Parker-Nance S, Davies-Coleman MT, Trindade M. (2017). Antibacterial Activities of Bacteria Isolated from the Marine Sponges Isodictya compressa and Higginsia bidentifera Collected from Algoa Bay, South Africa. Mar Drugs 15.

17. Van Soest RWM, Boury-Esnault N, Vacelet J, Dohrmann M, Erpenbeck D, De Voogd NJ, Santodomingo N, Vanhoorne B, Kelly M, Hooper JNA. (2012). Global diversity of sponges (Porifera). PloS one 7:e35105–e35105.

18. Mohamed M. Radwan SPM, Samir A. Ross. (2007). Two New Sulfated Sterols from the Marine Sponge Lendenfeldia dendyi NPC Natural Product Communications 2.

19. Liu Y, Liu R, Mao S-C, Morgan JB, Jekabsons MB, Zhou Y-D, Nagle DG. (2008). Molecular-Targeted Antitumor Agents. 19. Furospongolide from a Marine Lendenfeldia sp. Sponge Inhibits Hypoxia-Inducible Factor-1 Activation in Breast Tumor Cells. Journal of Natural Products 71:1854–1860.

20. Sera Y, Adachi K, Shizuri Y. (1999). A New Epidioxy Sterol as an Antifouling Substance from a Palauan Marine Sponge, Lendenfeldia chondrodes. Journal of Natural Products 62:152–154.

21. Thakur NL, Anil AC. (2000). Antibacterial Activity of the Sponge Ircinia Ramosa: Importance of its Surface-Associated Bacteria. Journal of Chemical Ecology 26:57–71.

22. Belma Konuklugġl BG. (2015). Antimicrobial activity of marine samples collected from the different coasts of turkey. Turkish Journal of Pharmaceutical Sciences 12.

23. Mohamed NM, Rao V, Hamann MT, Kelly M, Hill RT. (2008). Monitoring Bacterial Diversity of the Marine Sponge <em>Ircinia strobilina</em> upon Transfer into Aquaculture. Applied and Environmental Microbiology 74:4133.

24. Bassetti M, Baguneid M, Bouza E, Dryden M, Nathwani D, Wilcox M. (2014). European perspective and update on the management of complicated skin and soft tissue infections due to methicillin-resistant Staphylococcus aureus after more than 10 years of experience with linezolid. Clin Microbiol Infect 20 Suppl 4:3–18.

25. Dookie N, Rambaran S, Padayatchi N, Mahomed S, Naidoo K. (2018). Evolution of drug resistance in Mycobacterium tuberculosis: a review on the molecular determinants of resistance and implications for personalized care. Journal of Antimicrobial Chemotherapy 73:1138–1151.

26. Pourakbari B, Mamishi S, Mohammadzadeh M, Mahmoudi S. (2016). First-Line Anti-Tubercular Drug Resistance of Mycobacterium tuberculosis in IRAN: A Systematic Review. Frontiers in microbiology 7:1139–1139.

27. Lopez-Avalos G, Gonzalez-Palomar G, Lopez-Rodriguez M, Vazquez-Chacon CA, Mora-Aguilera G, Gonzalez-Barrios JA, Villanueva-Arias JC, Sandoval-Diaz M, Miranda-Hernández U, Alvarez-Maya I. (2017). Genetic diversity of Mycobacterium tuberculosis and transmission associated with first-line drug resistance: a first analysis in Jalisco, Mexico. Journal of Global Antimicrobial Resistance 11:90–97.

28. Evans-Illidge EA, Logan M, Doyle J, Fromont J, Battershill CN, Ericson G, Wolff CW, Muirhead A, Kearns P, Abdo D, Kininmonth S, Llewellyn L. (2013). Phylogeny drives large scale patterns in Australian marine bioactivity and provides a new chemical ecology rationale for future biodiscovery. PLoS One 8:e73800.

29. Simpson M, Poulsen S-A. (2014). An Overview of Australia’s Compound Management Facility: The Queensland Compound Library. ACS Chemical Biology 9:28–33.

30. Anonymous. (2018). Compounds Australia. https://www.griffith.edu.au/griffith-sciences/compounds-australia. Accessed 20 September 2018.

31. Bouslimani A, Sanchez LM, Garg N, Dorrestein PC. (2014). Mass spectrometry of natural products: current, emerging and future technologies. Natural Product Reports 31:718–729.

32. Kind T, Fiehn O. (2010). Advances in structure elucidation of small molecules using mass spectrometry. Bioanalytical reviews 2:23–60.

33. Herbert Júnior D, Nathalya Isabel de M, Antônio Eduardo Miller C. (2012). Electrospray Ionization Tandem Mass Spectrometry as a Tool for the Structural Elucidation and Dereplication of Natural Products: An Overview. In Prasain J (ed), Tandem Mass Spectrometry-Applications and Principles doi:10.5772/32680. InTech.

34. Pang S, Deng Y, Shi F. (2015). Synthesis of unsymmetric tertiary amines via alcohol amination. Chemical Communications 51:9471–9474.

35. Fujita K-i, Yamamoto K, Yamaguchi R. (2002). Oxidative Cyclization of Amino Alcohols Catalyzed by a Cp*Ir Complex. Synthesis of Indoles, 1,2,3,4-Tetrahydroquinolines, and 2,3,4,5-Tetrahydro-1-benzazepine. Organic Letters 4:2691–2694.

36. Cheng D, Wang Z, Xia Y, Wang Y, Zhang W, Zhu W. (2016). Catalytic amination of diethylene glycol with tertiarybutylamine over Ni-Al2O3 catalysts with different Ni/Al ratios. RSC Advances 6:102373–102380.

37. McCracken ST. (2010). Synthetic Studies of Biologically Active Natural Products – Ascididemin and 6-Substituted 2-Pyranones. Doctor of Philosophy Auckland, New Zealand.

38. Lindsay BS. (1998). Studies in Marine Natural Product Synthesis, lsolation and Ecology. Doctor of Philosophy. University of Auckland, University of Auckland.

39. Phuwapraisirisan P, Matsunaga S, van Soest RWM, Fusetani N. (2004). Shinsonefuran, a cytotoxic furanosesterterpene with a novel carbon skeleton, from the deep-sea sponge Stoeba extensa1See Ref. 1.1. Tetrahedron Letters 45:2125–2128.

40. Carr G. (2010). Bioactive Marine Natural Products: Isolation, Structure Elucidation And Synthesis Of Pharmacophore Analogues. Doctor of philosophy The university of british columbia Vancouver.

41. Veltri CA. (2009). Identification of cellular targets of marine natural products. Doctor of Philosophy The University of Utah United States.

42. Turkson NY. (2007). The Search for Efflux Pump Inhibitors from Marine Natural Products Doctor of Philosophy University of california, San diego.

43. Bontemps N, Bry D, Lopez-Legentil S, Simon-Levert A, Long C, Banaigs B. (2010). Structures and antimicrobial activities of pyridoacridine alkaloids isolated from different chromotypes of the ascidian Cystodytes dellechiajei. J Nat Prod 73:1044–8.

44. Dassonneville L, Wattez N, Baldeyrou B, Mahieu C, Lansiaux A, Banaigs B, Bonnard I, Bailly C. (2000). Inhibition of topoisomerase II by the marine alkaloid ascididemin and induction of apoptosis in leukemia cells. Biochem Pharmacol 60:527–37.

45. Kobayashi JC, J; Nakamura, H.; Ohizumi, Y.; Hirata, Y; Sasaki, T.; Ohta, T.; Nozoe, S. (1988). Tetrahedron Letters 29.

46. Schmitz FJdG, F. S.; Choi, y,; Hossain, M. B.; Rizvi, S. K.; van der Helm, D. (1990). Pure & Appl Chem 62.

47. Lindsay BSB, L. R.; Copp, B. R. (1995). Bio Med Chem Leff 5.

48. Bracher F. (1989). Heterocycles 29.

49. Moody CJR, C. W.; Thomas, R. (1990). Tetrahedron Letters 31.

50. Gellerman GR, A.; Kashman, Y. (1994). Synfhesis 3.

51. Bonnard I, Bontemps N, Lahmy S, Banaigs B, Combaut G, Francisco C, Colson P, Houssier C, Waring MJ, Bailly C. (1995). Binding to DNA and cytotoxic evaluation of ascididemin, the major alkaloid from the Mediterranean ascidian Cystodytes dellechiajei. Anticancer Drug Des 10:333–46.

52. Ralifo P, Sanchez L, Gassner NC, Tenney K, Lokey RS, Holman TR, Valeriote FA, Crews P. (2007). Pyrroloacridine alkaloids from Plakortis quasiamphiaster: structures and bioactivity. J Nat Prod 70:95–9.

53. Marshall KM, Andjelic CD, Tasdemir D, Concepción GP, Ireland CM, Barrows LR. (2009). Deoxyamphimedine, a Pyridoacridine Alkaloid, Damages DNA via the Production of Reactive Oxygen Species. Marine Drugs 7:196–209.

54. Bracher F. (1989). Total Synthesis of the Pentacyclic Alkaloid Ascididemin. Herrocycles 29: 2093–2095.

55. Banaigs SL-LNB-SXTB. (2007). Secondary metabolite and inorganic contents in Cystodytes sp. (Ascidiacea): temporal patterns and association with reproduction and growth. Mar Biol 151:293–299.

56. Kimura J, Ishizuka E, Nakao Y, Yoshida WY, Scheuer PJ, Kelly-Borges M. (1998). Isolation of 1-Methylherbipoline Salts of Halisulfate-1 and of Suvanine as Serine Protease Inhibitors from a Marine Sponge, Coscinoderma mathewsi. Journal of Natural Products 61:248–250.

57. Svendsen L, Blombäck B, Blombäck M, Olsson PI. (1972). Synthetic chromogenic substrates for determination of trypsin, thrombin and thrombin-like enzymes. Thrombosis Research 1:267–278.

58. Nakao Y, Matsunaga S, Fusetani N. (1995). Three more cyclotheonamides, C, D, and E, potent thrombin inhibitors from the marine sponge Theonella swinhoei. Bioorganic & Medicinal Chemistry 3:1115–1122.

59. Furuta A, Salam KA, Hermawan I, Akimitsu N, Tanaka J, Tani H, Yamashita A, Moriishi K, Nakakoshi M, Tsubuki M, Peng PW, Suzuki Y, Yamamoto N, Sekiguchi Y, Tsuneda S, Noda N. (2014). Identification and biochemical characterization of halisulfate 3 and suvanine as novel inhibitors of hepatitis C virus NS3 helicase from a marine sponge. Mar Drugs 12:462–76.

60. Cheuka PM, Mayoka G, Mutai P, Chibale K. (2016). The Role of Natural Products in Drug Discovery and Development against Neglected Tropical Diseases. Molecules 22:58.

61. Gordaliza M. (2009). Terpenyl-purines from the sea. Mar Drugs 7:833–49.

62. Giard DJ, Aaronson SA, Todaro GJ, Arnstein P, Kersey JH, Dosik H, Parks WP. (1973). In vitro cultivation of human tumors: establishment of cell lines derived from a series of solid tumors. J Natl Cancer Inst 51:1417–23.

63. Gaush CR, Hard WL, Smith TF. (1966). Characterization of an established line of canine kidney cells (MDCK). Proc Soc Exp Biol Med 122:931–5.

64. Tsuchiya S, Yamabe M, Yamaguchi Y, Kobayashi Y, Konno T, Tada K. (1980). Establishment and characterization of a human acute monocytic leukemia cell line (THP-1). Int J Cancer 26:171–6.

65. Aden DP, Fogel A, Plotkin S, Damjanov I, Knowles BB. (1979). Controlled synthesis of HBsAg in a differentiated human liver carcinoma-derived cell line. Nature 282:615–6.

66. Graham FL, Smiley J, Russell WC, Nairn R. (1977). Characteristics of a human cell line transformed by DNA from human adenovirus type 5. J Gen Virol 36:59–74.

67. Liberio MdS. (2014). Chemical and Biological Investigations of Anticancer Compounds from Australian Ascidians. B.Sc., M.Sc. Griffith University, Griffith University.

68. Lindsay BS. (1998). Studies in Marine Natural Product Synthesis, lsolation and Ecology.. PhD. University of Auckland, University of Auckland.

69. Poigny S, Nouri S, Chiaroni A, Guyot M, Samadi M. (2001). Total Synthesis and Determination of the Absolute Configuration of Coscinosulfate. A New Selective Inhibitor of Cdc25 Protein Phosphatase. The Journal of Organic Chemistry 66:7263–7269.

70. Bae J, Jeon J-e, Lee Y-J, Lee H-S, Sim CJ, Oh K-B, Shin J. (2011). Sesterterpenes from the Tropical Sponge Coscinoderma sp. Journal of Natural Products 74:1805–1811.

71. Kernan MR, Faulkner DJ. (1988). Sesterterpene sulfates from a sponge of the family Halichondriidae. The Journal of Organic Chemistry 53:4574–4578.

72. Loukaci A, Le Saout I, Samadi M, Leclerc S, Damiens E, Meijer L, Debitus C, Guyot M. (2001). Coscinosulfate, a CDC25 phosphatase inhibitor from the sponge Coscinoderma mathewsi. Bioorg Med Chem 9:3049–54.

